# Plant growth regulators ameliorate or exacerbate abiotic and biotic stress effects on Zea mays kernel weight in a genotype-specific manner

**DOI:** 10.1101/088195

**Authors:** Lauren Stutts, Yishi Wang, Ann E. Stapleton

## Abstract

Plant growth regulators have documented roles in plant responses to single stresses. In combined-stress environments, plants display novel genetic architecture for growth traits and the response to growth regulators is unclear. We investigated the role of plant growth regulators in combined-stress responses in Zea mays. Twelve maize inbreds were exposed to all combinations of the following stressors: drought, nitrogen, and density stress. Chemical treatments were utilized to alter balances of the hormones abscisic acid, gibberellic acid, and brassinosteroids. We found a significant difference between the seed weights of plants given different chemical treatments after accounting for differences in genotype and stress environments. We conclude that plant growth regulators have targets in combined-stress response pathways in Zea mays.

**HIGHLIGHT:** Plant growth regulators can ameliorate effects of combinations of abiotic and biotic stress in maize, in certain genotypes and under specific stress conditions.

## INTRODUCTION

Of all of the world’s grains, maize production is the largest by weight, and the United States is the top exporter of this grain (Capehart) USDA (2016). The large amounts of maize produced are used globally in various roles; maize remains a major source of food for both humans and animals around the world, and is also utilized in the production of biofuels. Maize production is continuously being improved by efforts in plant breeding and genetic modification, therefore maize genetics remains a central area of research. Geneticists aim to select for traits that will result in better protection against pests, more resistance to harsh environmental conditions, and as a more nutritious food source (Carena *et al.*, 2010). With the continued growth of the world’s population the agriculture industry faces a higher demand for grains and smaller land resources to meet this demand. Therefore, the overarching goal is to produce crops of a higher quality at higher quantity. Knowledge about plant response to stress at the molecular level is key to meeting higher demand, as a large proportion of crops are exposed to stress annually (Lobell and Gourdji, 2012).

### Response to Environmental Stress

The response of crops to abiotic and biotic stress has long been a focal point for agricultural research. A solid comprehension of the mechanisms plants use to combat environmental stresses such as drought, light-stress, fungal infections, and nutrient depletion, for example, allows researchers to develop plants that will be more resistant to these stresses (Taiz and Zeiger, 2006). Exposure of plants to stress at certain points of development can have detrimental impacts on growth and crop yield (Carena *et al.*, 2010). Significant decreases in corn grain yield and plant biomass can result from limitations in nitrogen availability, which is especially important in low-input smallholder settings (Weber *et al.*, 2012). Loss of plant biomass can also be seen in response to varying plant density, even in some modern maize hybrids (Tokatlidis *et al.*, 2011).

Source-sink balance is a key determinant of the final harvest weight of maize kernels, typically with an interaction between genotypes and environmental limits across years (Borrás *et al.*, 2004; Sala *et al.*, 2007; Boomsma *et al.*, 2009). Kernel weight is less affected by late abiotic stress than kernel number, and the kernel weight environmental response varied across hybrid genotypes (Slafer and Otegui, 2000). This makes kernel weight a useful trait across both basic research and applied agronomic experimentation (Kesavan *et al.*, 2013; Zhang *et al.*, 2016).

Increased attention has been given recently to plant responses to combined stresses (Rejeb *et al.*, 2014). Plant physiological responses to combined-stress are not additive; when exposed to two simultaneous stresses, portions of the two single-stress response genetic pathways are expressed, but not all (Mittler, 2006; Suzuki *et al.*, 2014). A synergistic response to drought and low nitrogen maize can be present and has been exploited for production via agronomic advice to reduce nitrogen fertilizer application under drought conditions (Bennett *et al.*, 1989; Weber *et al.*, 2012; Sadras and Richards, 2014), though this synergistic response is genetically variable and thus would not apply to all production settings. Beyond the nitrogen-drought interaction, additional non-linear combinatoric responses can be used to group maize genotypes into high and low input optimal types (Ruffo *et al.*, 2015).

A signaling network has been proposed by Makumburage et al. (Makumburage *et al.*, 2013), in which loci within individual stress response pathways repress loci in different stress response pathways. Evidence was provided that maize displays a novel genetic architecture in response to combined stress, relative to the architecture of genetic response to a single stress. Makumburage et al. (2013) observed that the interaction between two stress-response pathways in corn allowed improved growth under combined-stresses compared to what would be expected. Combining abiotic and biotic stressors, specifically plant density, also results in non-additive responses (Rossini *et al.*, 2011). Information about response to combined-stresses is relevant in the agriculture industry, because crops growing in the field encounter multiple stresses simultaneously throughout their life cycle, rather than only one stress in an otherwise controlled environment.

At the cellular level, it has been observed that once a plant has been exposed to one stress, the molecular response to a second stress can be altered (Rasmussen *et al.*, 2013). Furthermore, novel genes not expressed under either stress individually are expressed when the plant is exposed to both stresses simultaneously (Rizhsky *et al.*, 2004; Plessis *et al.*, 2015). Humbert et al. (Humbert *et al.*, 2013) presented data confirming that maize transcript-level response to drought varied depending on whether the plant was also under nitrogen stress.

### Genotypes

For this study, we selected a range of genotypes that were from temperate, tropical and mapping populations. The B73 and Mo17 inbreds are widely studied; B73 in particular was a key to improved germplasm in the single-cross hybrid era of maize breeding (Carena *et al.*, 2010). Improved tropical genotype CML103 was selected by CIMMYT breeders and is included in key diversity panels such as the NAM (McMullen *et al.*, 2009). We also chose a few genotypes from a widely used mapping population (more than 300 citations), which was derived from B73 and Mo17 with intermating to increase the number of recombination events (Lee *et al.*, 2002).

### Plant Hormones

Plant hormones have been long known to be mediators between the external environment and the internal activities of plants (Wilkinson *et al.*, 2012). Plant hormones regulate the growth and development of plants, stimulating seed germination, placement and growth of new organs, death and abscission of organs, ripening of fruit, and regulation of stomatal closure (Taiz and Zeiger, 2006). Hormones are involved in cross-talk between other pathways within the plant (Mittler *et al.*, 2011), and often play an integrator role between multiple pathways (Jaillais and Chory, 2010; Gómez-Cadenas *et al.*, 2014). Plant hormones are tightly intertwined with every aspect of the organism’s life. Due to their role as pathway integrators, we have focused on hormones as candidates for the interaction seen between stress-response pathways during multiple-stress responses.

Gibberellins are a group of plant hormones known to influence plant growth and development (Taiz and Zeiger, 2006). These compounds are synthesized in the chloroplasts, endoplasmic reticulum, and cytosol of plant cells, and transported via the xylem, and play a role in modulation of abiotic stress (Colebrook *et al.*, 2014). Abscisic acid has long been recognized for its role in plant response to water-limiting conditions. Abscisic acid is known to be a key player in regulating the opening and closing of stomata, by controlling the surrounding guard cells (Li *et al.*, 2006). There is evidence that abscisic acid may be active in helping plants to tolerate both short-term and long-term stress (Sreenivasulu *et al.*, 2012). Another group of plant signaling molecules known to have significant impacts on plant growth is brassinosteroids. These molecules are synthesized in the endoplasmic reticulum and bind neighboring cell surface receptors (Symons *et al.*, 2008), thereby initiating complex signaling pathways within the plant, allowing response to environmental conditions (Belkhadir and Chory, 2006).

### Plant Growth Regulators

Due to their effects on plant traits, hormones are targets of researchers attempting to influence these traits. “Plant Growth Regulator” is a term given to a large group of chemicals used to alter intrinsic levels of plant hormones. Many of these chemicals are sold commercially, and target the biosynthesis or degradation of plant hormones. In this study, we used three commercial plant growth regulators, along with direct application of gibberellic acid, to change hormone levels within individual plants. The three plant growth regulators used were paclobutrazol (PAC), uniconazole (UCN), and propiconazole (PCZ). These compounds are all triazoles, which target enzymes and result in inhibition of the synthesis of various compounds, such as GA, BR, and ABA. Some triazole compounds were originally used as fungicides (by limiting GA synthesis in fungi), and were later recognized for their effects on plant growth (Rademacher *et al.*, 1992). Paclobutrazol is commonly used to limit stem elongation in crops. The compound inhibits synthesis of gibberellic acid by preventing formation of the precursor molecule kaurenoic acid (Hedden and Graebe, 1985). The limitation of stem elongation in crops is beneficial because it prevents lodging, or stem breakage, when top-heavy plants are exposed to adverse conditions. Uniconazole is another regulator used to limit plant height. Uniconazole has also been shown to increase drought tolerance in Arabidopsis thaliana (Saito *et al.*, 2006). These effects are achieved by inhibiting synthesis of GA and BR, and inhibiting the breakdown of ABA. Treatment of maize with propiconazole also results in dwarf phenotypes, via inhibition of BR synthesis (Hartwig *et al.*, 2012).

In this study we investigated the role of hormones in plant responses to combined stresses, via manipulation of intrinsic hormone levels of plants grown in single-stress and combined-stress environments. We predicted that an alteration in hormone balance would alter combined-stress response pathways and ultimately alter phenotypic response. We found that for certain genotypes, a creation of hormone imbalance within the plants altered the seed weight trait response to combined-stress environments.

## MATERIALS AND METHODS

### Co-localization of Combined-stress QTL and Hormone-responsive Genes

Combined stress quantitative trait loci (QTL) were determined from previous field experiments (Stapleton, personal communication; (Makumburage *et al.*, 2013). These QTL are regions of the maize genome determined to be important to certain plant traits in response to various stressors in the field. QTL were identified under either combined drought and UV stress or combined drought and nitrogen stress. Regions of the maize genome were identified as significant contributors to phenotypic traits such as plant height and root biomass. The coordinates of these combined-stress regions were identified by using the Locus Lookup function on MaizeGDB (http://www.maizegdb.org/) to find marker positions on the maize genome. The coordinates of these regions were placed into Qteller (http://www.qteller.com/), which returned a list of all genes present in the multiple-stress QTL. This gene list was then used in agriGO, which used Fisher’s significance test (α=0.05) to determine what lists of genes compose the combined-stress QTL gene set.

To determine which genes compose hormone pathways in maize, a list of genes known to be responsive to hormones was compiled from information obtained in the literature and from MaizeGDB. The physical locations of hormone-responsive genes on the maize genome were determined using MaizeGDB (RefGen v2 sequence map). The number of hormone-responsive genes co-localized with combined-stress QTL was determined, and the ComBin function in Microsoft Excel was used to evaluate the significance of genes falling into these QTL, by calculating the combinatorial probability that they would be in those locations by chance (Balint-Kurti *et al.*, 2010).

### Field Design and Implementation of Abiotic Stress Conditions

Plants were grown in an experimental plot at the Central Crops Research Station in Clayton, North Carolina, Latitude 35.66979°, Longitude-78.4926° from April 12 to August 30, 2013. The field was arranged in a strip plot design, in which the plants were exposed to up to three of the following stresses: nitrogen deprivation, drought, and high density stress. The field was divided into eight sections as shown in Figure 1, and each of the sections received a combination of between zero and three of the stresses previously mentioned, so that all possible stress combinations were included. Drought stress was imposed by lack of irrigation to stressed sections. Water was supplied to irrigated portions as needed during silking and grain fill. A nitrogen-stressed environment was created by lack of nitrogen application. Other nutrient-containing fertilizers were applied equally across all sections of the field, in accordance with standard maize growth practice at this site. Density stress was implemented during planting, with seeds planted four inches apart in stressed sections, rather than eight inches apart as was done in control sections.

**Figure 1:**
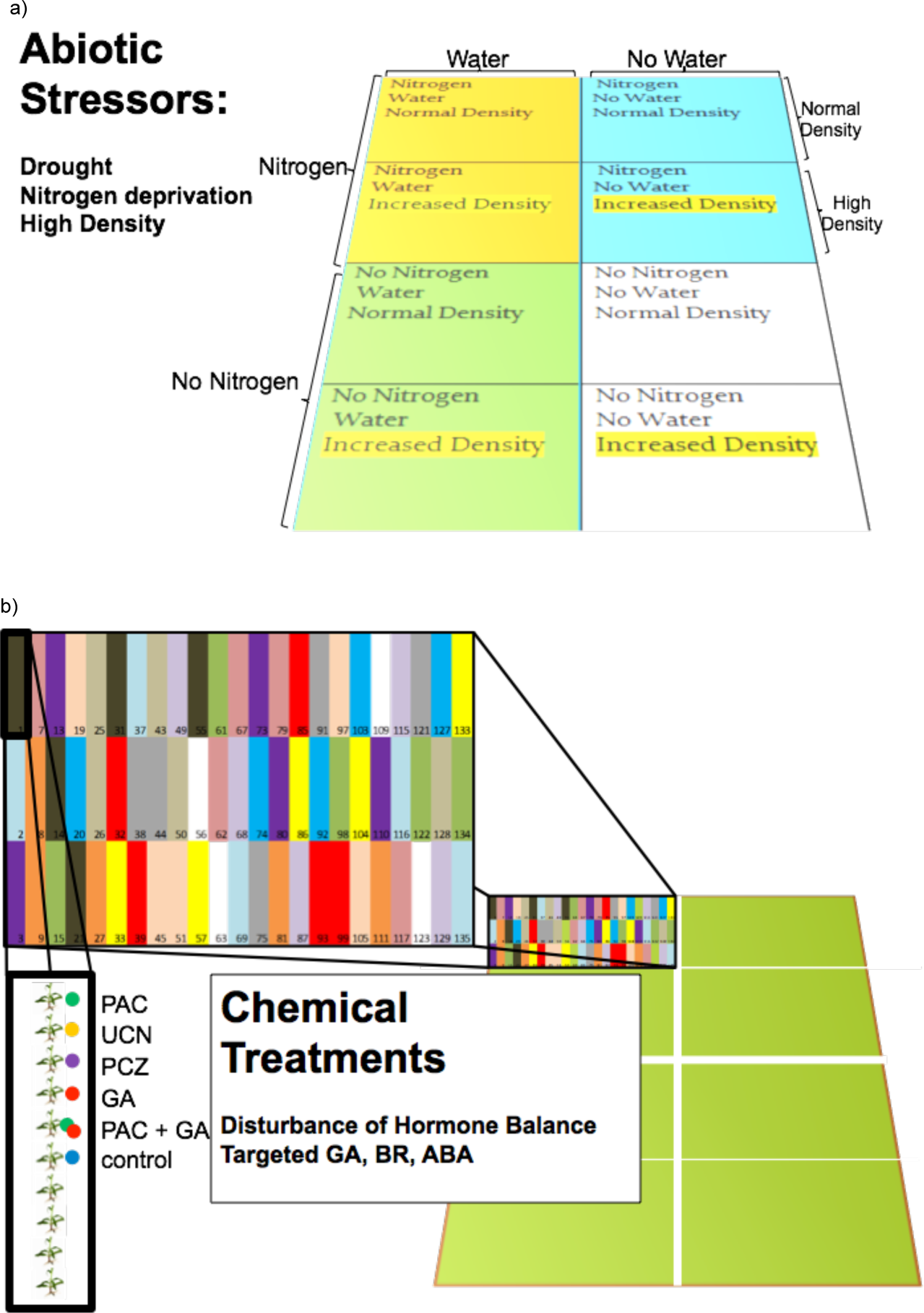
Figure 1 Field layout, with eight stress sections (Fig. 1a), twelve genotypes replicated five times in each stress section (colored bars in Fig. 1b), and six hormone treatments given in each genotype section (colored dots in Fig. 1b).

### Genotypes

Twelve genotypes of Zea mays were used in this study; these genotypes had previously been observed to have differences in response to stress (A. Stapleton, personal communication, (Makumburage and Stapleton, 2011; Makumburage *et al.*, 2013) for the trait plant height. Seeds of inbreds B73 and Mo17 and the IBM94RIL population were obtained from the maize co-op (http://maizecoop.cropsci.uiuc.edu/). Inbreds Oh7b, Oh43, CML103 and LH132 were kindly provided by Dr. James Holland, USDA & NCSU. Six recombinant inbred lines from the IBM (intermated B73 and Mo17) mapping population were utilized, as well as their parent lines, B73 and Mo17 (Lee *et al.*, 2002). All lines in the IBM population have differing combinations of alleles from these parent genotypes, which confer varying levels of tolerance to abiotic stressors. We used three genotypes previously classified as stress intolerant (Mo298, Mo352, and Mo360), and three lines with a prior identification as higher tolerance for stress (Mo017, Mo276, and Mo287). The inbred lines LH132, CML103, Oh7B, and Oh43, which were also previously shown to have differing responses to combinations of drought and low nitrogen (A. Stapleton, personal communication), were included as well. As seen in Figure 1, all twelve Zea mays genotypes were represented within each of the eight stress environments. The Oh43 genotype plants germinated poorly and several plots were lost during the experiment, so that genotype was removed from further analysis. Genotypes of the RIL lines were confirmed by SNP analysis performed by a commercial genotyping service (RapidGenomics, Inc, Gainesville, FL, USA).

### Chemical Treatment for the Creation of Hormone Imbalances

Chemical treatments were used to interrupt either synthesis or degradation of target hormones. To affect balance between the hormones gibberellic acid, brassinosteroids, and abscisic acid, we used the following chemicals: paclobutrazol (PAC), uniconazole (UCN), propiconazole (PCZ), and gibberellic acid (GA). One of six chemical treatment types was given to individual plants within each environment: PAC, UCN, PCZ, GA, PAC with GA, and a control treatment with no chemical application, as shown in Figure 2. Ten mL of a 50 ppm concentration of each chemical was applied to each treated plant. For plants given the PAC and GA treatment, 10 mL of both solutions were applied to give a total of 20 ml per whorl. Solutions were pipetted directly into the whorls of plants during the 4-6 leaf stage of development, five weeks after planting. Solutions were prepared from commercial plant growth regulators. Piccolo ornamental plant growth regulator (manufactured by Fine Agrochemicals Ltd, Walnut Creek, CA, USA), which has a 4000 ppm concentration of paclobutrazol, was diluted to yield a 50 ppm paclobutrazol solution. Propiconazole 14.3 (Quali-Pro, Pasadena, TX, USA) contains a 143,000 ppm concentration of propiconazole, and was diluted to yield a 50 ppm solution. Sumagic plant growth regulator (Valent Biosciences, Libertyville, IL, USA) contains a 550 ppm concentration of uniconazole, which was diluted to yield a 50 ppm uniconazole solution. 99% gibberellic acid (Acros Organics, Thermo Fisher, Pittsburgh, PA, USA) was dissolved in water to yield a 50 ppm GA solution.

**Figure 2:**
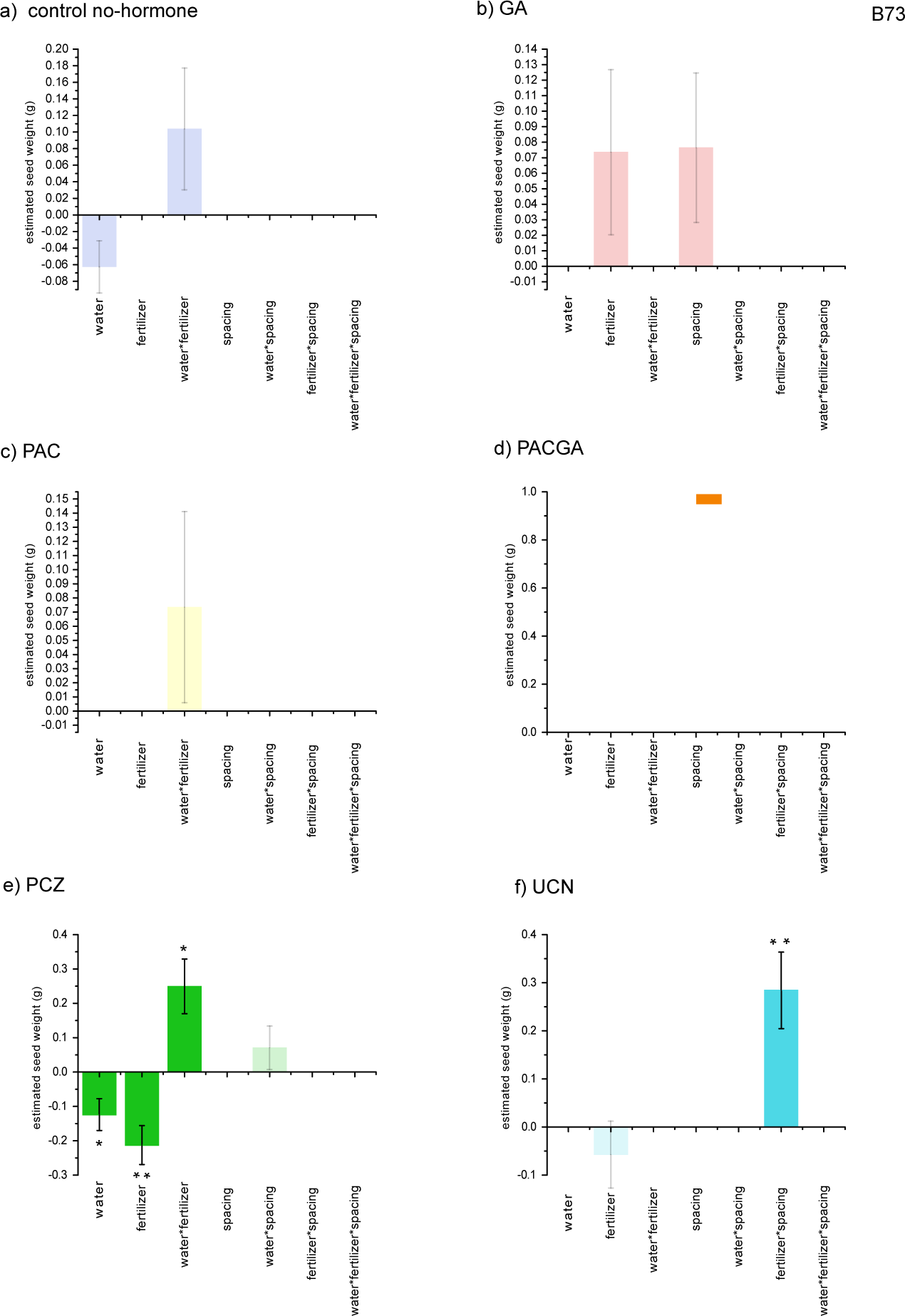
Figure 2 Lasso seed weight effect estimates on the original parameter scale (colored bars) and standard errors (black lines within bars) are plotted for each plant growth regulator and abiotic-biotic stress combination in genotype B73. Estimates shrunk to zero are not listed, which non-significant estimates are indicated as faded-color bars. Estimates significant after multiple–comparison Bonferroni correction within lines are indicated with single asterisks (*) and estimates significant after experiment-wise correction are indicated with two asterisks (**). a) Lasso parameter estimates for the control plants that were not treated with plant growth regulators. b) Lasso parameter estimates for plants treated with gibberellic acid (GA). c) Lasso parameter estimates for plants treated with paclobutrazol (PAC). d) Lasso parameter estimates for plants treated with both GA and PAC simultaneously. e) Lasso parameter estimates for plants treated with propiconazole (PCZ). f) Lasso parameter estimates for plants treated with uniconazole (UCN).

### Data Collection and Analysis

Ear traits were measured after completion of reproductive development and drydown. The uppermost ear was harvested from each plant, and cob diameter (in millimeters) and 20-seed weights were collected from each ear. Seed weight was recorded as the collective weight (in grams) of 20 kernels removed from the center of each cob. If 20 kernels were not present, the seed weight was recorded as zero. The seed weight data are included as supplemental file 1, with metadata descriptions as supplemental file 2. A factorial model with hormone treatment, stress treatments, and all interaction terms for each genotype was used to compare environment, genotype, and hormone treatment group means for the traits measured. Models were fit by generalized regression using a zero-inflated gamma distribution, with Adaptive Lasso and AICc, using SAS JMP v12 (SAS Inc, Cary, NC). First, the control no-hormone treatment was analyzed for each stress factor separately as a full factorial for each line, then the growth regulator treatments were analyzed; significance thresholds were adjusted to an experiment-wise Bonnferroni multiple-test corrected threshold of P < 0.005.

The pattern of seed weight response to stress for each plant growth regulator was analyzed using a novel non-parametric method. A detailed description of the new method, with simulation results, power analysis and theoretical foundations, is provided at https://arxiv.org/abs/1611.04619. Briefly, we use bootstrap sampling to compare all pairwise subsamples and determine whether a stress combination’s values are higher, lower or equal. This allows us to detect differences in the trend of the response between two plant growth regulators; multiple pairwise comparisons then identify significant contrasts between treatments. The method was designed for short-series data with zero-inflated observations.

## RESULTS

### QTL Overlaps

The set of chromosomal loci for hormone-related genes and the loci identified as important for multiple-stress plant traits overlapped significantly (P=0.001). A overview map of the loci is available in supplemental Figure 1.

### Factorial Seed Weight Responses to Stress and Plant Growth Regulators

The overall pattern of reduced seed weight was non-linear, with the best-fit distribution a spline rather than a normal distribution; test of linear fit gave an R-squared of 0.17 as compared to the spline-fit R-squared of 0.34 (data not shown). We note that, overall, the seed weight of the drought low-nitrogen treatment was higher than the full-water low nitrogen treatment, indicating that these stresses combined in a nonlinear way in our experiment. Genotypes CML103 and Mo360 exhibited significantly reduced seed weight when exposed to increased abiotic and biotic stress and stress combinations without growth regulator (Fig. 4a, Fig. 11a, Supplemental Results File 1 for factor estimates and all P-values); all other genotypes exhibited no significant effect of abiotic-biotic stress on seed weight. Plant height, tassel architecture and cob diameter traits show the same general trends as seed weight (Stutts, 2014).

For each genotype we analyzed the effect of plant growth regulators and stress factors (Figures 2-11). The effect of stress and growth regulator was specific to the genotype examined, so we discuss each genotype individually below; for some genotypes there were both positive and negative effects on seed weight in some stress-chemical combinations while other genotypes had more uniform responses to chemical treatment.

#### B73

In the B73 genotype (Figure 2) we saw significant increases and decreases in seed weight with plant growth regulator treatment. Specifically, PCZ increased the sensitivity of this inbred to drought and low nitrogen, decreasing seed weight, but increased the seed weight in the combined drought-low nitrogen plot (Fig. 2e). This low-nitrogen amelioration of drought yield depression is a common response pattern in commercial genotypes. The UCN treatment in B73 generated an increased seed weight only in the low-nitrogen-fertilizer and dense spacing plot (Fig. 2f).

#### Mo17

The Mo17 inbred genotype seed weight was not strongly affected by abiotic and biotic stress (Fig. 3a). Application of PAC exaggerates the effect of low nitrogen and high density applied together (Fig. 3c), resulting in significantly lower seed weights. This PAC-sensitivity was no longer detectable when PAC and GA were applied together (Fig. 3d). The effect of combining low-nitrogen conditions, drought, and PAC plus GA resulted in significantly increased seed weights (Fig. 3d), which were not apparent in the individual stress plots or plant growth regulator treatments (Fig. 3b, c).

**Figure 3:**
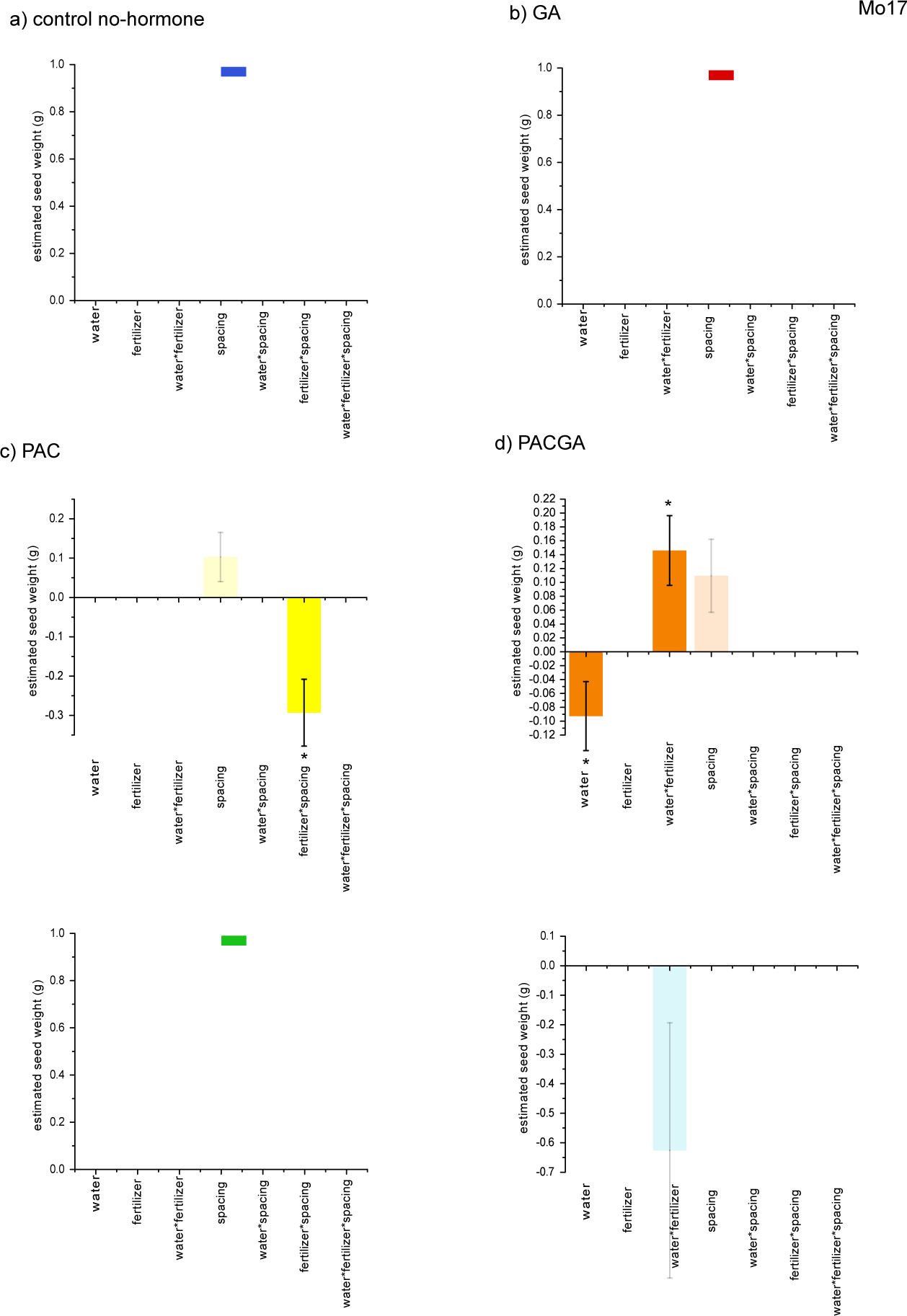
Figure 3 Lasso seed weight effect estimates on the original parameter scale (colored bars) and standard errors (black lines within bars) are plotted for each plant growth regulator and abiotic-biotic stress combination in genotype Mo17. Estimates shrunk to zero are not listed, which non-significant estimates are indicated as faded-color bars. Estimates significant after multiple–comparison Bonferroni correction within lines are indicated with single asterisks (*) and estimates significant after experiment-wise correction are indicated with two asterisks (**). a) Lasso parameter estimates for the control plants that were not treated with plant growth regulators. b) Lasso parameter estimates for plants treated with gibberellic acid (GA). c) Lasso parameter estimates for plants treated with paclobutrazol (PAC). d) Lasso parameter estimates for plants treated with both GA and PAC simultaneously. e) Lasso parameter estimates for plants treated with propiconazole (PCZ). f) Lasso parameter estimates for plants treated with uniconazole (UCN).

#### CML103

Decreases in seed weight under low nitrogen conditions were observed in CML103 (Fig. 4a) and those decreases were unaffected by GA or PAC (Fig. 4b, c). However, there was no significant decrease in the other plant growth regulator treatments, likely due to an increase in the variance of seed weight that is visible and longer standard error bars in Fig. 4d, e and f.

**Figure 4:**
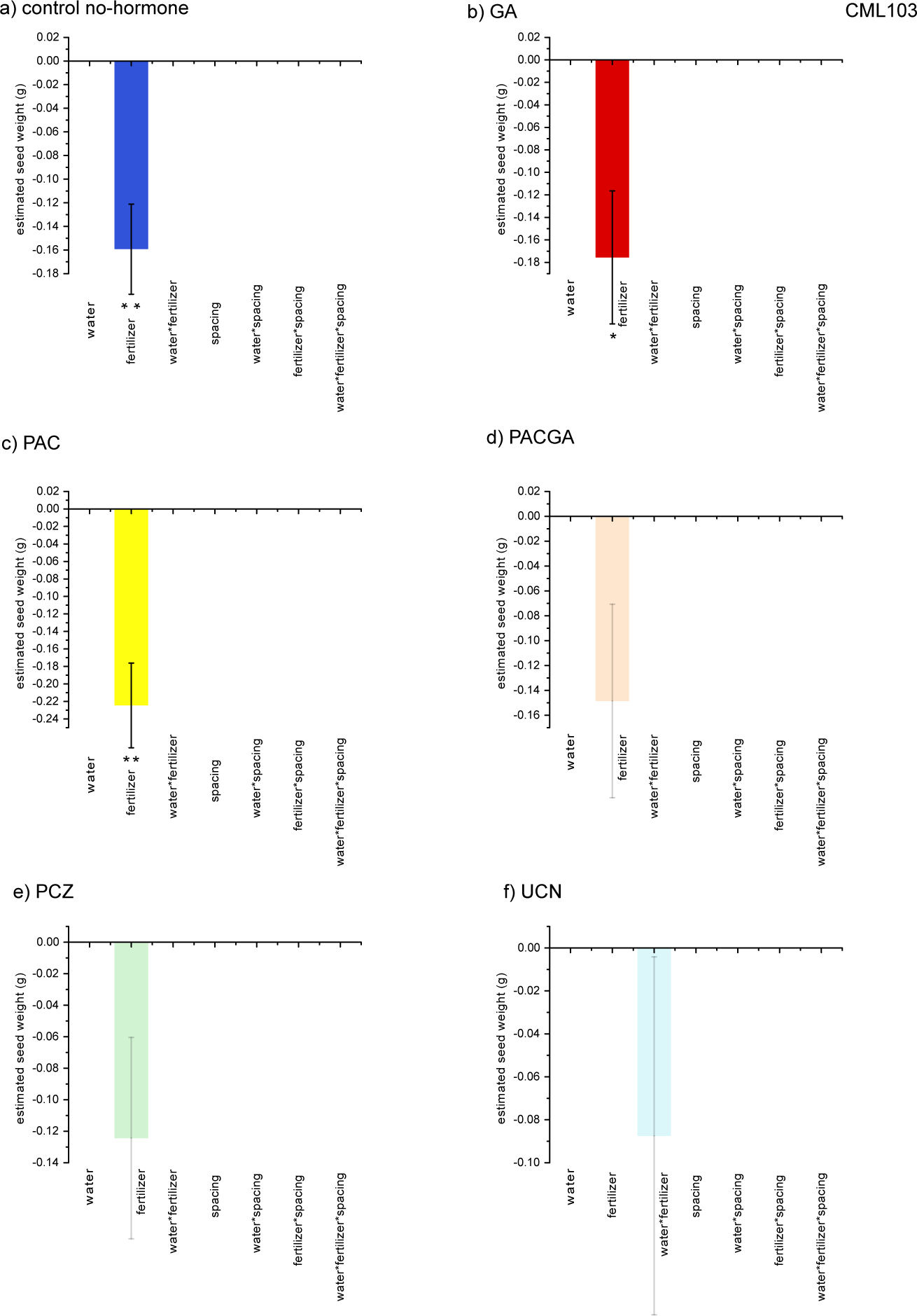
Figure 4 Lasso seed weight effect estimates on the original parameter scale (colored bars) and standard errors (black lines within bars) are plotted for each plant growth regulator and abiotic-biotic stress combination in genotype CML103. Estimates shrunk to zero are not listed, which non-significant estimates are indicated as faded-color bars. Estimates significant after multiple–comparison Bonferroni correction within lines are indicated with single asterisks (*) and estimates significant after experiment-wise correction are indicated with two asterisks (**). a) Lasso parameter estimates for the control plants that were not treated with plant growth regulators. b) Lasso parameter estimates for plants treated with gibberellic acid (GA). c) Lasso parameter estimates for plants treated with paclobutrazol (PAC). d) Lasso parameter estimates for plants treated with both GA and PAC simultaneously. e) Lasso parameter estimates for plants treated with propiconazole (PCZ). f) Lasso parameter estimates for plants treated with uniconazole (UCN).

#### Oh7b

Inbred Oh7B shows no significant reduction in seed weight under abiotic and biotic stress (fig. 5a), grouping this genotype with B73 and Mo17 and contrasting these three genotypes with the low-nitrogen sensitivity of CML103. In the GA treatment, Oh7b seed weights declined in low nitrogen conditions. This decline was not observed in PAC (Fig. 4c), but was still present in PACGA (Fig. 5d). Thus, in this genotype the sensitivity to low nitrogen conferred by GA is independent. Treatment with PCZ also exaggerates the decline in seed weight under low nitrogen (Fig. 4e). There was a density-PAC interaction, with PAC treatment conferring increased sensitivity to high plant density and density under drought (Fig. 4c).

**Figure 5:**
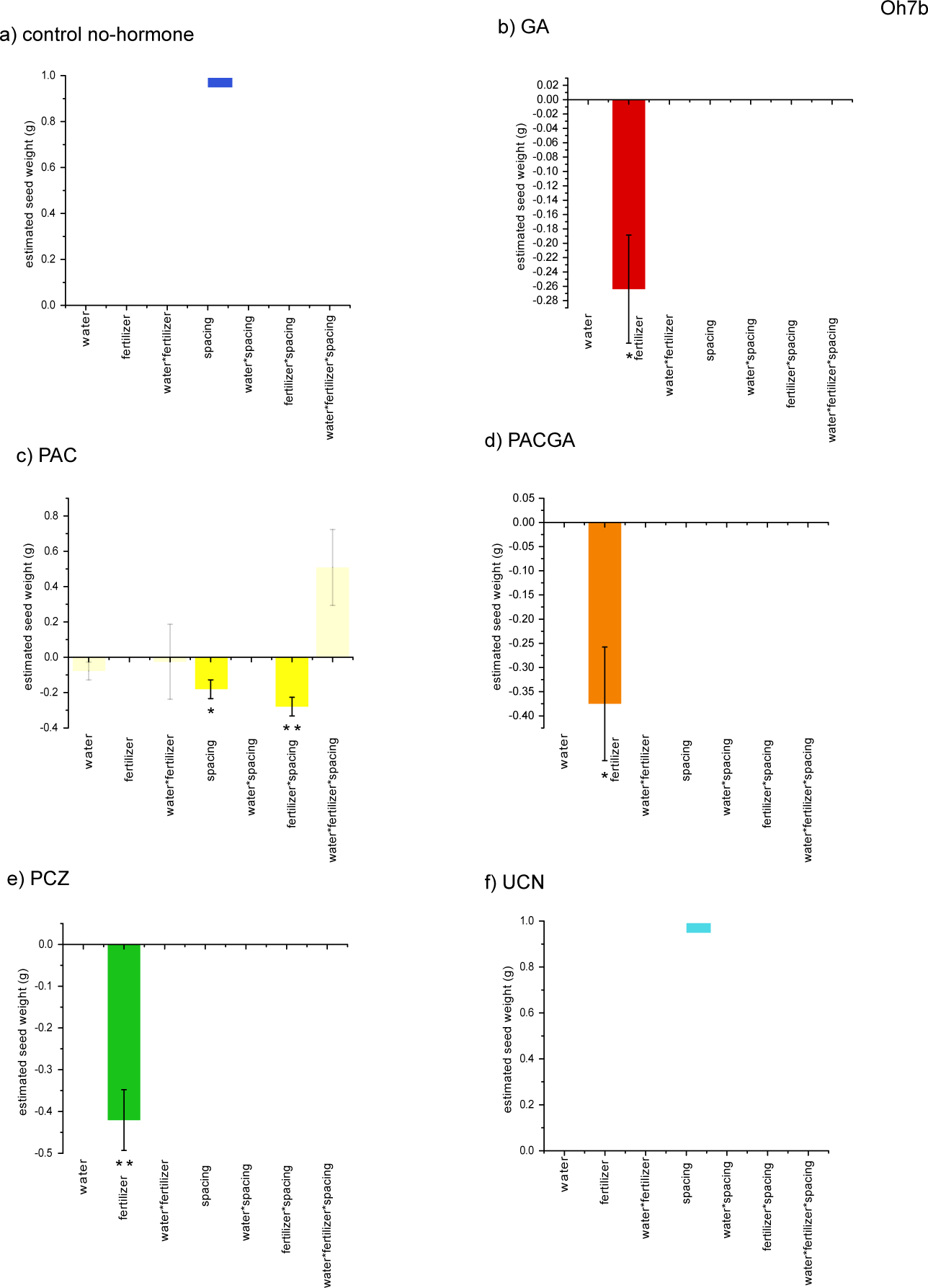
Figure 5 Lasso seed weight effect estimates on the original parameter scale (colored bars) and standard errors (black lines within bars) are plotted for each plant growth regulator and abiotic-biotic stress combination in genotype Oh7b. Estimates shrunk to zero are not listed, which non-significant estimates are indicated as faded-color bars. Estimates significant after multiple–comparison Bonferroni correction within lines are indicated with single asterisks (*) and estimates significant after experiment-wise correction are indicated with two asterisks (**). a) Lasso parameter estimates for the control plants that were not treated with plant growth regulators. b) Lasso parameter estimates for plants treated with gibberellic acid (GA). c) Lasso parameter estimates for plants treated with paclobutrazol (PAC). d) Lasso parameter estimates for plants treated with both GA and PAC simultaneously. e) Lasso parameter estimates for plants treated with propiconazole (PCZ). f) Lasso parameter estimates for plants treated with uniconazole (UCN).

#### LH132

In the ex-plant-variety-protection inbred LH132 the effect of low nitrogen was most prominent, with significant decline in seed weight in the low-nitrogen PAC and low-nitrogen PCZ treatments (Fig. 6c, e). A dependence on nitrogen input might be expected for an inbred selected for good performance under high-input modern agronomic conditions. There is no significant drought or combined stress effect on seed weight in any plant growth regulator treatment, again supporting a history of selection for insensitivity to common uncontrolled environmental conditions and the new agronomic norm of high plant density in this genotype.

**Figure 6:**
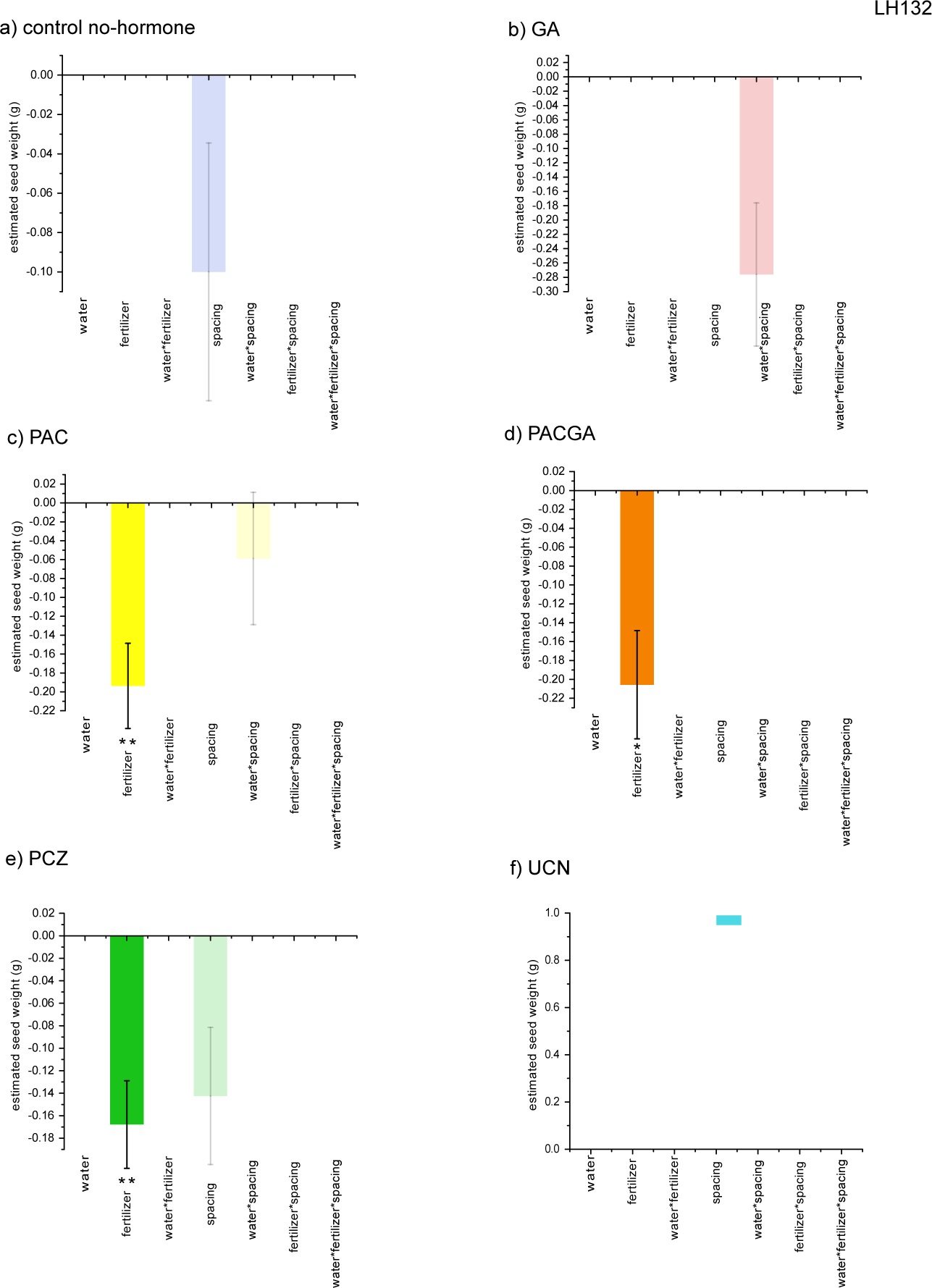
Figure 6 Lasso seed weight effect estimates on the original parameter scale (colored bars) and standard errors (black lines within bars) are plotted for each plant growth regulator and abiotic-biotic stress combination in genotype LH132. Estimates shrunk to zero are not listed, which non-significant estimates are indicated as faded-color bars. Estimates significant after multiple–comparison Bonferroni correction within lines are indicated with single asterisks (*) and estimates significant after experiment-wise correction are indicated with two asterisks (**). a) Lasso parameter estimates for the control plants that were not treated with plant growth regulators. b) Lasso parameter estimates for plants treated with gibberellic acid (GA). c) Lasso parameter estimates for plants treated with paclobutrazol (PAC). d) Lasso parameter estimates for plants treated with both GA and PAC simultaneously. e) Lasso parameter estimates for plants treated with propiconazole (PCZ). f) Lasso parameter estimates for plants treated with uniconazole (UCN).

#### Mo017

In the IBM RIL Mo017 the no-plant-growth-regulator exhibited no change in seed weight across stress treatments (Fig. 7a). The PAC treatment conferred additional resistance to drought, however, with a significant increase in seed weight (Fig. 7c). A seed weight increase was not detectable in GA or PACGA treatments, suggesting that GA over-rides the beneficial effect of PAC under drought conditions.

**Figure 7:**
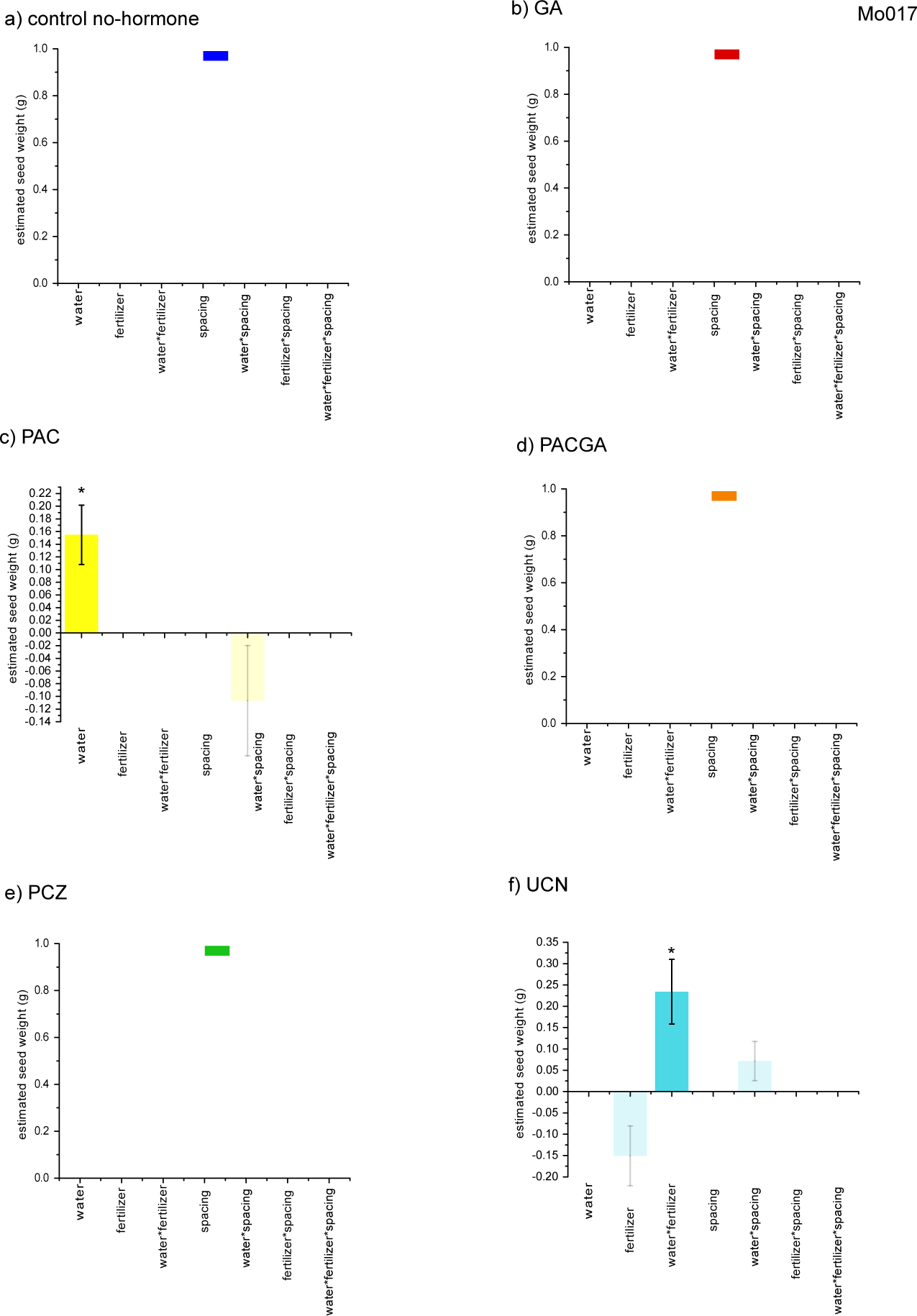
Figure 7 Lasso seed weight effect estimates on the original parameter scale (colored bars) and standard errors (black lines within bars) are plotted for each plant growth regulator and abiotic-biotic stress combination in genotype IBMRIL Mo017. Estimates shrunk to zero are not listed, which non-significant estimates are indicated as faded-color bars. Estimates significant after multiple–comparison Bonferroni correction within lines are indicated with single asterisks (*) and estimates significant after experiment-wise correction are indicated with two asterisks (**). a) Lasso parameter estimates for the control plants that were not treated with plant growth regulators. b) Lasso parameter estimates for plants treated with gibberellic acid (GA). c) Lasso parameter estimates for plants treated with paclobutrazol (PAC). d) Lasso parameter estimates for plants treated with both GA and PAC simultaneously. e) Lasso parameter estimates for plants treated with propiconazole (PCZ). f) Lasso parameter estimates for plants treated with uniconazole (UCN).

#### Mo0276

RIL Mo276 exhibited an increased seed weight in the triple-stress condition when GA was applied (Fig. 8b). The PAC treatment resulted in lower seed weight in low nitrogen (Fig. 8c). In the PACGA combined treatment, both the positive GA effect and the negative PAC effect were lost, thus suggesting that the PAC and GA pathways interact in this genotype.

**Figure 8:**
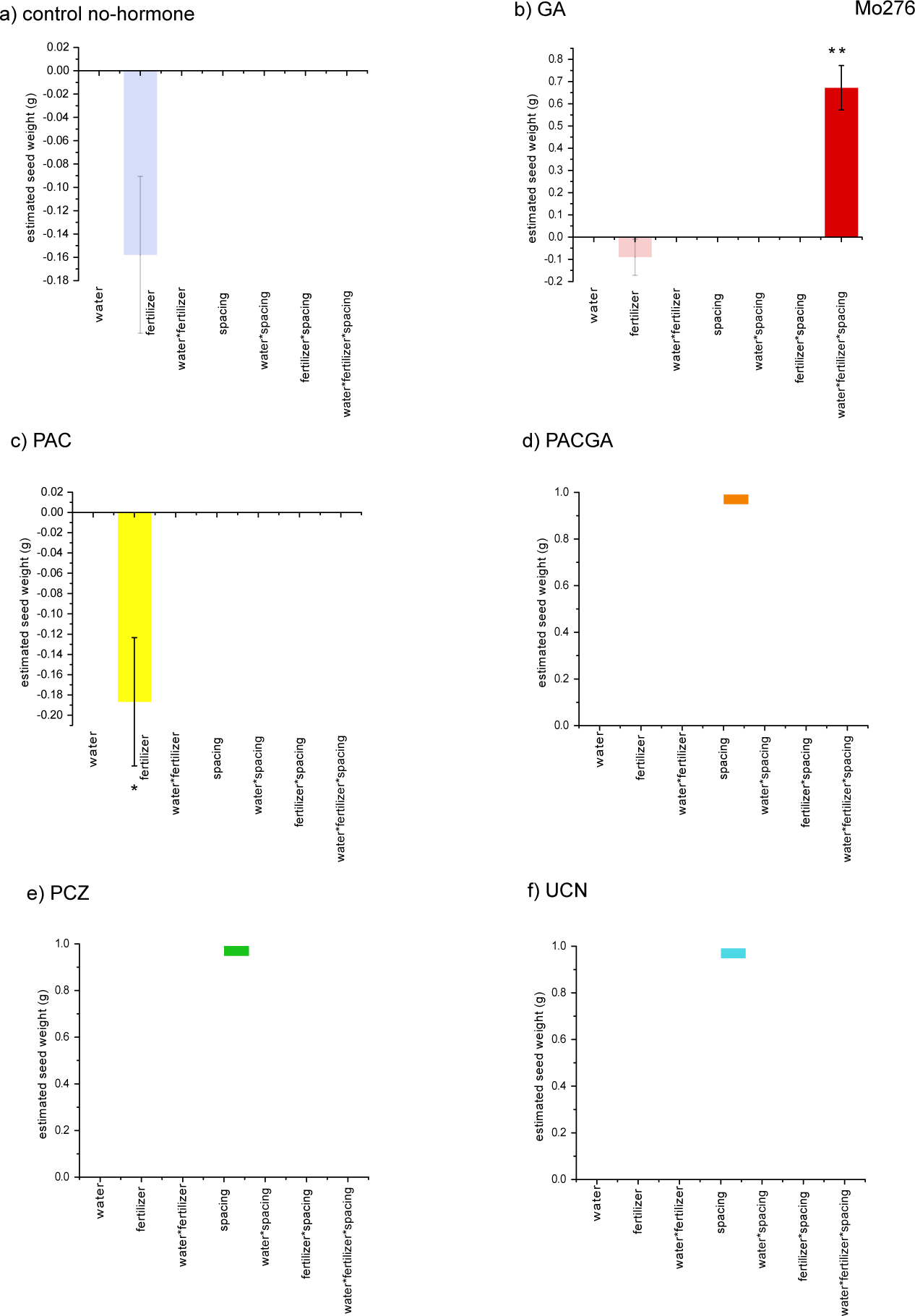
Figure 8 Lasso seed weight effect estimates on the original parameter scale (colored bars) and standard errors (black lines within bars) are plotted for each plant growth regulator and abiotic-biotic stress combination in genotype IBMRIL Mo276. Estimates shrunk to zero are not listed, which non-significant estimates are indicated as faded-color bars. Estimates significant after multiple–comparison Bonferroni correction within lines are indicated with single asterisks (*) and estimates significant after experiment-wise correction are indicated with two asterisks (**). a) Lasso parameter estimates for the control plants that were not treated with plant growth regulators. b) Lasso parameter estimates for plants treated with gibberellic acid (GA). c) Lasso parameter estimates for plants treated with paclobutrazol (PAC). d) Lasso parameter estimates for plants treated with both GA and PAC simultaneously. e) Lasso parameter estimates for plants treated with propiconazole (PCZ). f) Lasso parameter estimates for plants treated with uniconazole (UCN).

#### Mo287

In the Mo287 RIL, the PAC treatment resulted in decreased seed weight in drought and low-nitrogen conditions, with a synergistic increase in seed weight in combined drought and low-nitrogen(Fig. 9c). This genotype plus PAC thus recreate a typical agronomic nonlinear pattern of stress response.

**Figure 9:**
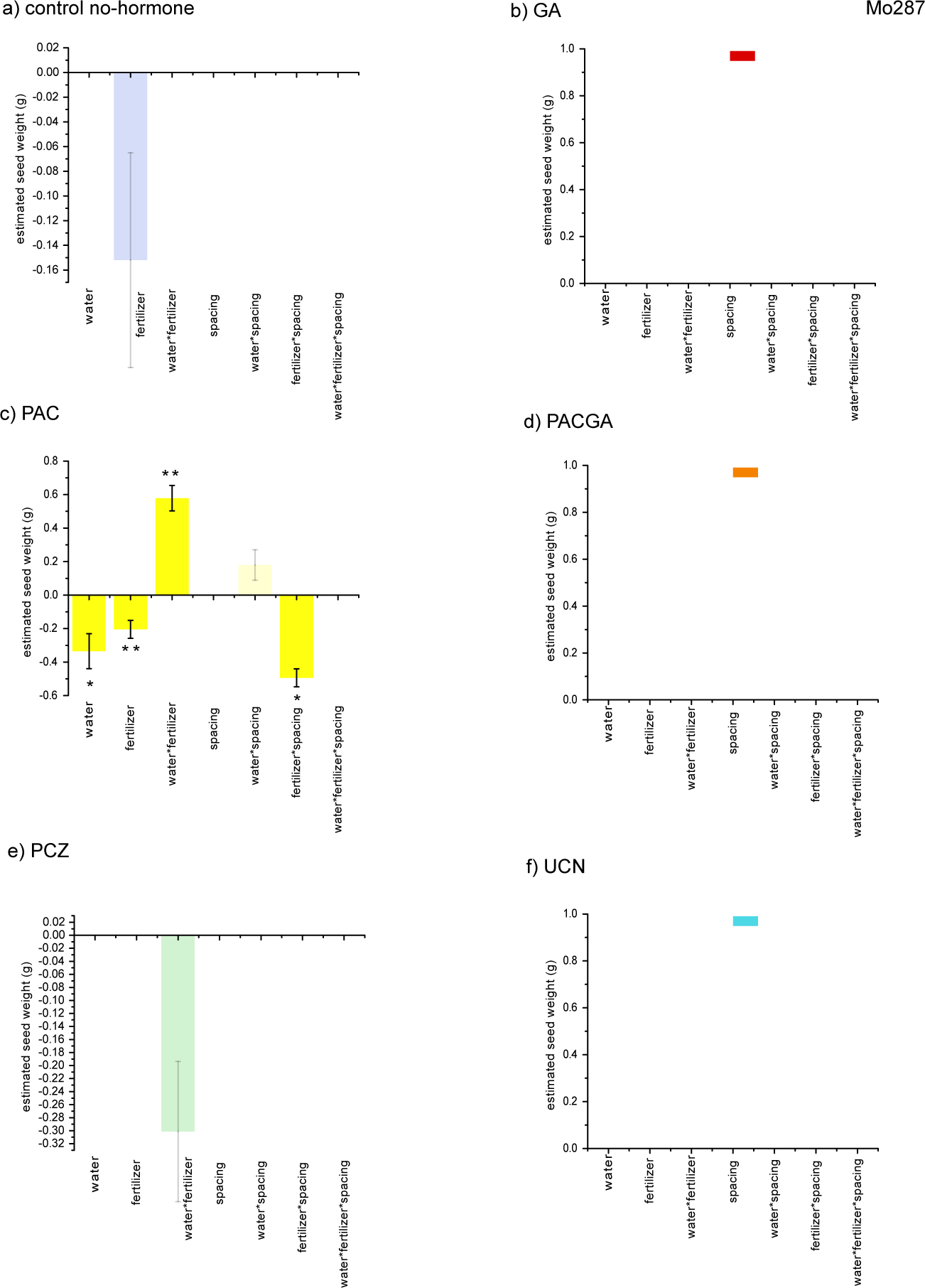
Figure 9 Lasso seed weight effect estimates on the original parameter scale (colored bars) and standard errors (black lines within bars) are plotted for each plant growth regulator and abiotic-biotic stress combination in genotype IBMRIL Mo287. Estimates shrunk to zero are not listed, which non-significant estimates are indicated as faded-color bars. Estimates significant after multiple–comparison Bonferroni correction within lines are indicated with single asterisks (*) and estimates significant after experiment-wise correction are indicated with two asterisks (**). a) Lasso parameter estimates for the control plants that were not treated with plant growth regulators. b) Lasso parameter estimates for plants treated with gibberellic acid (GA). c) Lasso parameter estimates for plants treated with paclobutrazol (PAC). d) Lasso parameter estimates for plants treated with both GA and PAC simultaneously. e) Lasso parameter estimates for plants treated with propiconazole (PCZ). f) Lasso parameter estimates for plants treated with uniconazole (UCN).

#### Mo352

In IBM94 RIL Mo352 (Fig. 10) low nitrogen has a more deleterious effect on seed weight in the presence of PAC, PACGA, and PCZ (Fig 10 c, d, and e). Drought also conferred lower seed weight in the PCZ-treated plants (Fig. 10e).

**Figure 10:**
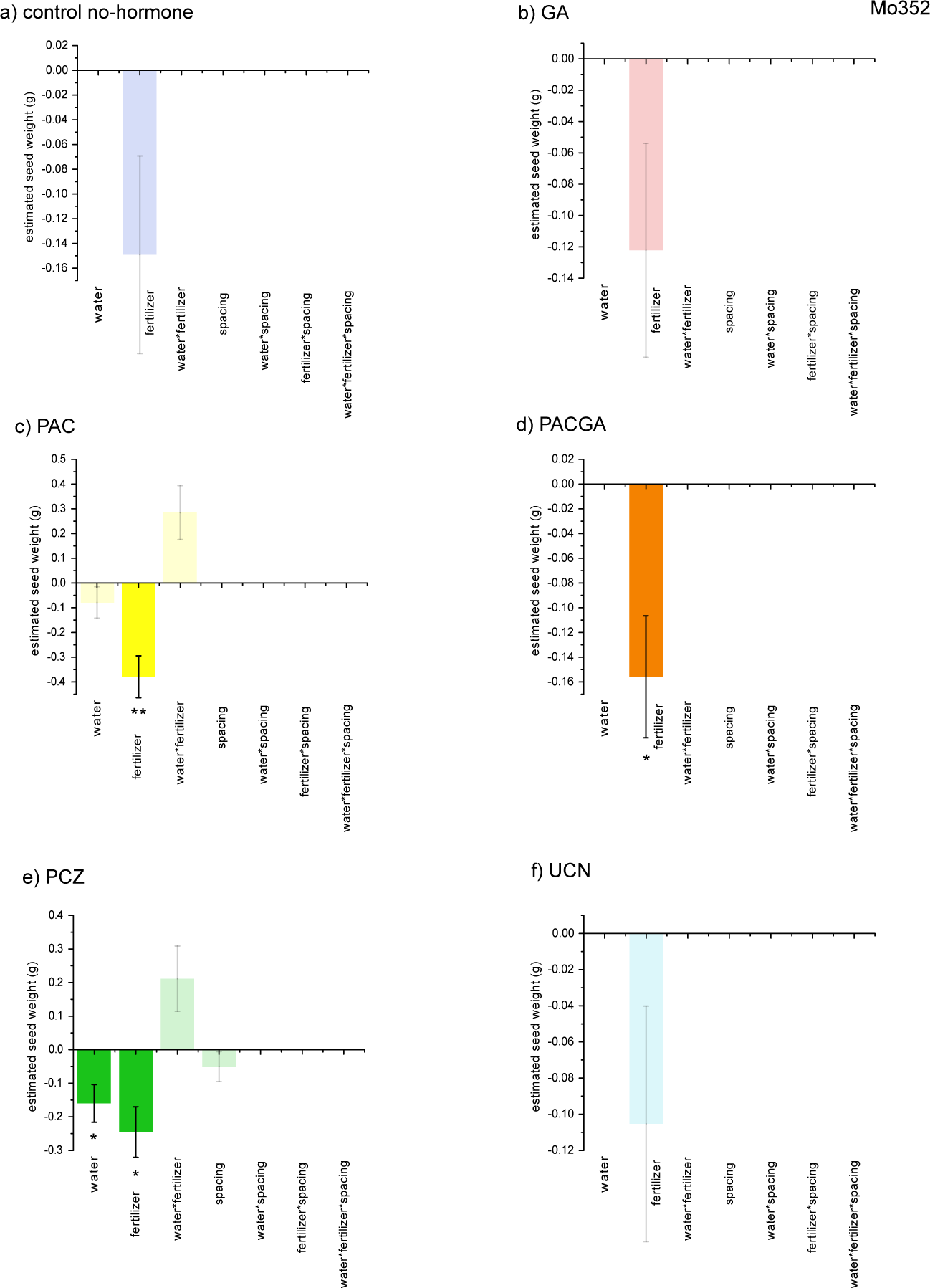
Figure 10 Lasso seed weight effect estimates on the original parameter scale (colored bars) and standard errors (black lines within bars) are plotted for each plant growth regulator and abiotic-biotic stress combination in genotype IBMRIL Mo352. Estimates shrunk to zero are not listed, which non-significant estimates are indicated as faded-color bars. Estimates significant after multiple–comparison Bonferroni correction within lines are indicated with single asterisks (*) and estimates significant after experiment-wise correction are indicated with two asterisks (**). a) Lasso parameter estimates for the control plants that were not treated with plant growth regulators. b) Lasso parameter estimates for plants treated with gibberellic acid (GA). c) Lasso parameter estimates for plants treated with paclobutrazol (PAC). d) Lasso parameter estimates for plants treated with both GA and PAC simultaneously. e) Lasso parameter estimates for plants treated with propiconazole (PCZ). f) Lasso parameter estimates for plants treated with uniconazole (UCN).

#### Mo360

In RIL Mo360, the control no-hormone stress treatments exhibited a higher seed weight at high density and a lower seed weight at high density with drought (Fig. 11a). This effect was not observed in any of the plant growth regulator treatments (Fig 11b-f), suggesting that regulator treatment reduced stress effects on seed weight. The PACGA treatment resulted in a lower seed weight in the low-nitrogen and high density stress setting (Fig. 11d).

**Figure 11:**
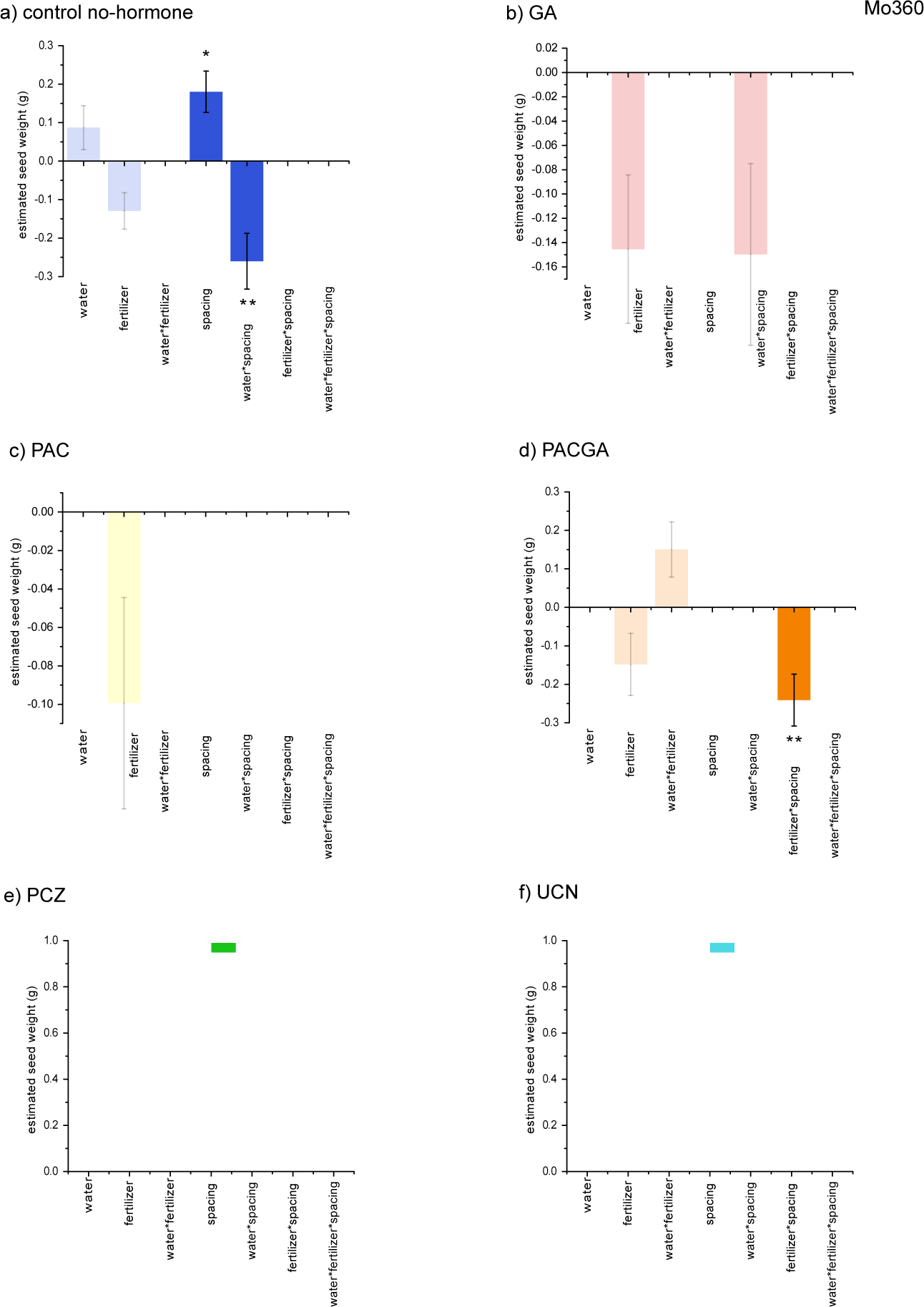
Figure 11 Lasso seed weight effect estimates on the original parameter scale (colored bars) and standard errors (black lines within bars) are plotted for each plant growth regulator and abiotic-biotic stress combination in genotype IBMRIL Mo360. Estimates shrunk to zero are not listed, which non-significant estimates are indicated as faded-color bars. Estimates significant after multiple–comparison Bonferroni correction within lines are indicated with single asterisks (*) and estimates significant after experiment-wise correction are indicated with two asterisks (**). a) Lasso parameter estimates for the control plants that were not treated with plant growth regulators. b) Lasso parameter estimates for plants treated with gibberellic acid (GA). c) Lasso parameter estimates for plants treated with paclobutrazol (PAC). d) Lasso parameter estimates for plants treated with both GA and PAC simultaneously. e) Lasso parameter estimates for plants treated with propiconazole (PCZ). f) Lasso parameter estimates for plants treated with uniconazole (UCN).

### Genotype-Regulator Combinations with Positive Effects on Seed Weight

While the majority of the significant stress effects on seed weight were negative, with lower seeds weights in the presence of drought and low nitrogen, there were a few combination with increased seed mass. Relative to the control plants with no hormone treatment, some growth regulators were able increase seed weights. In B73 both UCN and PCZ conferred increased seed weight in combined stress conditions (Fig. 1e and f). For inbred Mo17, PAC plus GA treatment had significantly higher seed weights in the low-nitrogen and drought combined stress condition (Fig. 2d). In four of the six RILs, there were positive estimated effects; specifically, Mo017 PAC in drought (Fig. 7c), Mo276 GA triple stress (Fig. 8b), Mo287 PAC low-nitrogen plus drought (Fig. 9c), and finally, Mo360 high density with no growth regulator (Fig. 11a).

### Comparisons Across Genotypes

Comparison of experiment-wise significant effect directions across all genotypes (Fig. 2-11 two-asterisk bars) resulted in a set of twelve significantly lowered seed weights and three increased seed weight effects. PAC contributed to both increased and decreased seed weight interactions across all genotypes, while GA and UCN were only found to have positive effects on seed weights. PCZ effects were always in the direction of decreased seed weight. Comparison of B73 and Mo17 parental inbreds to RILs suggested that a variety of effects could segregate into offspring and thus the genetic architecture of the response may be complex. There was one experiment-wise-significant effect (a decline in seed weight in the PAC plus low-nitrogen combination) that was found in two of the RILs-Mo287 and Mo352.

### Stress Pattern Differences

As our main interest is in the comparison of the pattern of plant growth regulator effects on seed weight under abiotic and biotic stress, we developed (cite Yishi’s proof paper here) and applied a novel all-pairwise non-parametric method for comparison of trends in seed weight across stress levels and between growth regulator treatments. We found six significant differences in the pattern of weight change as increasing levels of stress and plant growth regulators were applied (Fig. 12). In Fig. 12, stress levels are indicated with low levels on the left of the graph x-axes, to contrast with our factorial analysis (Figs. 2-11). Inbred B73 exhibited significant differences in seed weight in the GA growth regulator treatment as compared to the PACGA treatment (Fig. 12a), with the PACGA treatment conferring a more sigmoidal pattern with a stronger contrast between high and low nitrogen levels.

The next five trend differences (Fig. 12 b-f) were more complex than B73, with both increased and decreased seed weight across the combinations. Each genotype with a significant effect had a different pattern of response (Fig. 12 b-f). The PAC and PACGA treatments were significantly different in seed weight in both Mo276 and Mo298 (Fig. 12 c, e), though the pattern of response was shaped differently in those two genotypes. CML103 (Fig. 12 b) and Mo352 (Fig 12f) exhibited significant differences in pattern with PCZ; there was no significant trend in any genotype with a UCN treatment.

This nonparametric trend analysis provides a complementary view of data trends when compared to our factorial LASSO analysis. For example, trend analysis indicated that PAC and PACGA were different in Mo298, while there were no significant effects detected in this genotype in the factorial LASSO analysis. The trend analysis highlights nonlinear responses that may be of use in planning selection schemes; for example, in CML103 the PCZ treatment ameliorated the loss of seed weight under high stress relative to the GA treatment, at the cost of overall equivalent seed weight under optimum conditions (Fig. 12b). In the Mo352 genotype, the PACGA treatment gave maximum seed weights under regular density and low nitrogen, while in high density and low nitrogen the PCZ treatment exhibited higher kernel weight (Fig. 12f). Another example of such a crossover interaction is in the Mo287 genotype, with GA treatment showing a crossover interaction with control no-hormone when low-nitrogen regular spacing full water is compared to standard-nitrogen regular spacing full water (Fig. 12d). Comparison of trends in this way might allow custom plant growth regulator recommendations that incorporate weather trends.

DISCUSSION

As expected, we observed significant differences between plant trait means in different stress environments, with relatively small effects of abiotic and density stress on seed weight and more substantial effects on plant height (Stutts, 2014). High density planting has been shown to increase biomass for certain traits in corn (Murphy *et al.*, 1996), which is consistent with our observations of plant height (Stutts, 2014). The mid-range effects of abiotic stress on seed weights provided us with the ability to detect both exaggerated abiotic and combination stress effects with plant growth regulator treatment and reduced, ameliorating effects of the chemical application. We used prior information on plant heights in nitrogen and drought to classify genotypes for inclusion in our experiment, and all but one were consistent with their prior classification, even though we changed to use of seed weights as the focal trait.

Our observation of the specificity of genotype-stress-plant growth regulator interactions is consistent with results using a different experimental design and commercial hybrid genotypes (Ruffo *et al.*, 2015). In the Ruffo et al. work commercial hybrids were classified into two groups, those responsive to high inputs and those with uniform performance across a range of inputs. The high-input-optimal group equivalent in our experiment would be genotype B73, which exhibited a strong positive response to plant growth regulator treatment in several plant growth regulator-stress environments. Differences in plant traits were observed between group means for genotypes, as expected. In our study we focused on differences within genotypes, across different stress combinations. For example, the group of genotypes that fared significantly better in the control environment did not consistently respond optimally in other environments; these are consistent with the Ruffo et al. 2015 ‘uniform’ genotype group. This reflects results in other studies, which have reported a significant interaction between genotype and environment (Nzuve *et al.*, 2013). A significant interaction between genotype and environment informs us that the effects of genotype are not consistent across all levels of environmental stress, and that in return, stress combination does not have identical effects on different genotypes.

We observed significant differences between trait means of plants given differing hormone treatments. Plant growth regulator treatment in some cases ameliorated the effects of combined stress on plants. It was predicted that exogenous hormone application would affect plant response to stress combination, as it has been shown that abiotic stressors cause changes in endogenous hormone levels in maize (Pirasteh-Anosheh *et al.*, 2013) and plant growth regulators are in common use in production settings. The effects of hormone treatment were not, however, identical in different stress environments in this study. The resultant phenotypic changes varied between positive and negative, depending on stress combination. For example, exogenous gibberellic acid treatment did not have the same effects on nitrogen-stressed plants that it did on plants in other environments.

The results of this study indicate that plant hormones do play a role in response to combined stresses. Furthermore, it shows us that hormone balance does not have consistent effects on phenotype across all levels of genotype and environment. When targeting plant hormone balance, we must also consider genotype and environment to predict plant growth effects. Beneficial future work would include investigation into how hormone balance affects combined-stress response at the molecular level. The discovery and manipulation of hormone-responsive genes important in various stress combinations could have implications for the plant breeding industry.

## SUPPLEMENTARY DATA

**Supplemental Figure 1 multiple-stress QTL results.png** Map of QTL bin positions and overlap of hormone-related *Zea mays* genes.

**Supplemental File 1 EnvTreat seedweight zeros.csv** Seed weight data.

**Supplemental File 2METADATA for EnvTreat seedeight zeros.txt** Column header, unit and cell information for interpretation of Supplemental File 1 data.

**Supplemental ResultsFile1a controlnohormone byLine stressfactorial.docx** Full output from LASSO analysis for the no-hormone treatment.

**Supplemental ResultsFile1b GAonly byLine lasso stressfactorial.docx** Full output from lasso analysis for the gibberellic acid treatment.

**Supplemental ResultsFile1c PAConly lasso byline stressfactorial.docx** Full output from lasso analysis for the pacbutrazol treatment.

**Supplemental ResultsFile1d PCZlasso byline stressfactorial.docx** Full output from lasso analysis for the propiconazole treatment.

**Supplemental ResultsFile1e PACGAlasso byline stressfactorial.docx** Full output from lasso analysis for the no-hormone treatment.

**Supplemental ResultsFile1f UCNlasso byline stressfullfactorial.docx** Full output from lasso analysis for the uniconazole treatment.

**Supplemental Table1 nonparametric pattern output.docx** Full output from the R code for our novel non-parametric trend analysis.

## ACKNOWLEDGMENTS

We appreciate data analysis assistance from Dr. Susan Simmons and seed processing assistance from Stapleton lab members. This project was partially supported by the National Research Initiative Competitive Grant no. 2009-35100-05028 from the USDA National Institute of Food and Agriculture.

## LITERATURE CITED

Balint-Kurti P, Simmons SJ, Blum JE, Ballare CL, Stapleton AE. 2010. Maize Leaf Epiphytic Bacteria Diversity Patterns Are Genetically Correlated with Resistance to Fungal Pathogen Infection. Molecular Plant-Microbe Interactions 23, 473–484.

Belkhadir Y, Chory J. 2006. Brassinosteroid signaling: a paradigm for steroid hormone signaling from the cell surface. Science (New York, N.Y.) 314, 1410–1411.

Bennett JM, Mutti LSM, Rao PSC, Jones JW. 1989. Interactive effects of nitrogen and water stresses on biomass accumulation, nitrogen uptake, and seed yield of maize. Field Crops Research 19, 297–311.

Boomsma CR, Santini JB, Tollenaar M, Vyn TJ. 2009. Maize Morphophysiological Responses to Intense Crowding and Low Nitrogen Availability: An Analysis and Review. Agronomy Journal 101, 1426.

Borrás L, Slafer GA, Otegui ME. 2004. Seed dry weight response to source–sink manipulations in wheat, maize and soybean: a quantitative reappraisal. Field Crops Research 86, 131–146.

Capehart T. USDA Economic Research Service – Corn.

Carena MJ, Hallauer AR, Miranda Filho JB, Filho JBM. 2010. Quantitative Genetics in Maize Breeding. Springer New York.

Colebrook EH, Thomas SG, Phillips AL, Hedden P. 2014. The role of gibberellin signalling in plant responses to abiotic stress. The Journal of Experimental Biology 217, 67–75.

Gómez-Cadenas A, Ollas C de, Manzi M, Arbona V. 2014. Phytohormonal Crosstalk Under Abiotic Stress. In: Tran, L-SP, In: Pal S, eds. Phytohormones: A Window to Metabolism, Signaling and Biotechnological Applications. Springer New York, 289–321.

Hartwig T, Corvalan C, Best NB, Budka JS, Zhu J-Y, Choe S, Schulz B. 2012. Propiconazole Is a Specific and Accessible Brassinosteroid (BR) Biosynthesis Inhibitor for Arabidopsis and Maize. PLOS ONE 7, e36625.

Hedden P, Graebe JE. 1985. Inhibition of gibberellin biosynthesis by paclobutrazol in cell-free homogenates ofCucurbita maxima endosperm andMalus pumila embryos. Journal of Plant Growth Regulation 4, 111.

Humbert S, Subedi S, Cohn J, Zeng B, Bi Y-M, Chen X, Zhu T, McNicholas PD, Rothstein SJ. 2013. Genome-wide expression profiling of maize in response to individual and combined water and nitrogen stresses. BMC genomics 14, 3.

Jaillais Y, Chory J. 2010. Unraveling the paradoxes of plant hormone signaling integration. Nature Structural & Molecular Biology 17, 642–645.

Kesavan M, Song JT, Seo HS. 2013. Seed size: a priority trait in cereal crops. Physiologia Plantarum 147, 113–120.

Lee M-LT, Sharopova N, Beavis WD, Grant D, Katt M, Blair D, Hallauer A. 2002. Expanding the genetic map of maize with the intermated B73 x Mo17 (IBM) population. Plant Molecular Biology 48, 453–461.

Li S, Assmann SM, Albert R. 2006. Predicting Essential Components of Signal Transduction Networks: A Dynamic Model of Guard Cell Abscisic Acid Signaling. PLoS Biol 4, 10.

Lobell DB, Gourdji SM. 2012. The Influence of Climate Change on Global Crop Productivity. Plant Physiology 160, 1686–1697.

Makumburage GB, Richbourg HL, LaTorre KD, Capps A, Chen C, Stapleton AE. 2013. Genotype to phenotype maps: multiple input abiotic signals combine to produce growth effects via attenuating signaling interactions in maize. G3: Genes | Genomes | Genetics 3, 2195–2204.

Makumburage GB, Stapleton AE. 2011. Phenotype uniformity in combined-stress environments has a different genetic architecture than in single-stress treatments. Frontiers in Plant Science 2.

McMullen MD, Kresovich S, Villeda HS, et al. 2009. Genetic Properties of the Maize Nested Association Mapping Population. Science 325, 737–740.

Mittler R. 2006. Abiotic stress, the field environment and stress combination. Trends in Plant Science 11, 15–19.

Mittler R, Vanderauwera S, Suzuki N, Miller G, Tognetti VB, Vandepoele K, Gollery M, Shulaev V, Van Breusegem F. 2011. ROS signaling: the new wave? Trends in plant science 16, 300–309.

Murphy SD, Yakubu Y, Weise SF, Swanton CJ. 1996. Effect of planting patterns and inter-row cultivation on competition between corn (Zea mays) and late emerging weeds. Weed science.

Nzuve F, Githiri S, Mukunya DM, Gethi J. 2013. Analysis of Genotype x Environment Interaction for Grain Yield in Maize Hybrids. Journal of Agricultural Science 5, 75.

Pirasteh-Anosheh H, Emam Y, Pessarakli M. 2013. Changes in Endogenous Hormonal Status in Corn (zea Mays) Hybrids Under Drought Stress. Journal of Plant Nutrition 36, 1695–1707.

Plessis A, Hafemeister C, Wilkins O, Gonzaga ZJ, Meyer RS, Pires I, Mu¨ller C, Septiningsih EM, Bonneau R, Purugganan M. 2015. Multiple abiotic stimuli are integrated in the regulation of rice gene expression under field conditions. eLife 4, e08411.

Rademacher W, Temple-smith KE, Griggs DL, Hedden P. 1992. The mode of action of acylcyclohexanediones — a new type of growth retardant. In: Karssen, CM, In: van Loon, LC, In: Vreugdenhil D, eds. Current Plant Science and Biotechnology in Agriculture. Progress in Plant Growth Regulation. Springer Netherlands, 571–577.

Rasmussen S, Barah P, Suarez-Rodriguez MC, Bressendorff S, Friis P, Costantino P, Bones AM, Nielsen HB, Mundy J. 2013. Transcriptome responses to combinations of stresses in Arabidopsis. Plant Physiology 161, 1783–1794.

Rejeb IB, Pastor V, Mauch-Mani B. 2014. Plant Responses to Simultaneous Biotic and Abiotic Stress: Molecular Mechanisms. Plants 3, 458–475.

Rizhsky L, Liang H, Shuman J, Shulaev V, Davletova S, Mittler R. 2004. When Defense Pathways Collide. The Response of Arabidopsis to a Combination of Drought and Heat Stress. Plant Physiol. 134, 1683–1696.

Rossini MA, Maddonni GA, Otegui ME. 2011. Inter-plant competition for resources in maize crops grown under contrasting nitrogen supply and density: Variability in plant and ear growth. Field Crops Research In Press, Corrected Proof.

Ruffo ML, Gentry LF, Henninger AS, Seebauer JR, Below FE. 2015. Evaluating Management Factor Contributions to Reduce Corn Yield Gaps. Agronomy Journal 107, 495.

Sadras VO, Richards RA. 2014. Improvement of crop yield in dry environments: benchmarks, levels of organisation and the role of nitrogen. Journal of Experimental Botany 65, 1981–1995.

Saito S, Okamoto M, Shinoda S, et al. 2006. A plant growth retardant, uniconazole, is a potent inhibitor of ABA catabolism in Arabidopsis. Bioscience, Biotechnology, and Biochemistry 70, 1731–1739.

Sala RG, Westgate ME, Andrade FH. 2007. Source/sink ratio and the relationship between maximum water content, maximum volume, and final dry weight of maize kernels. Field Crops Research 101, 19–25.

Slafer GA, Otegui ME. 2000. Physiological Bases for Maize Improvement. CRC Press.

Sreenivasulu N, Harshavardhan VT, Govind G, Seiler C, Kohli A. 2012. Contrapuntal role of ABA: does it mediate stress tolerance or plant growth retardation under long-term drought stress? Gene 506, 265–273.

Stutts L. 2014. Hormones modify combined-stress responses in Zea mays. M.S. thesis, University of North Carolina Wilmington.

Suzuki N, Rivero RM, Shulaev V, Blumwald E, Mittler R. 2014. Abiotic and biotic stress combinations. New Phytologist 203, 32–43.

Symons GM, Ross JJ, Jager CE, Reid JB. 2008. Brassinosteroid transport. Journal of Experimental Botany 59, 17–24.

Taiz L, Zeiger E. 2006. Plant Physiology. Sunderland, MA: Sinauer Associates.

Tokatlidis I, Haş V, Mylonas I, Haş I, Evgenidis G, Melidis V, Copandean A, Ninou E. 2011. Density effects on environmental variance and expected response to selection in maize (Zea mays L.). Euphytica 174, 283–291.

Weber VS, Melchinger AE, Magorokosho C, Makumbi D, BÃ¤nziger M, Atlin GN. 2012. Efficiency of Managed-Stress Screening of Elite Maize Hybrids under Drought and Low Nitrogen for Yield under Rainfed Conditions in Southern Africa. Crop Sci. 52, 1011–1020.

Wilkinson S, Kudoyarova GR, Veselov DS, Arkhipova TN, Davies WJ. 2012. Plant hormone interactions: innovative targets for crop breeding and management. Journal of Experimental Botany 63, 3499–3509.

Zhang X, Hirsch CN, Sekhon RS, Leon N de, Kaeppler SM. 2016. Evidence for maternal control of seed size in maize from phenotypic and transcriptional analysis. Journal of Experimental Botany 67, 1907–1917.

